# Auxin coreceptor IAA17/AXR3 controls cell elongation in *Arabidopsis thaliana* root by modulation of auxin and gibberellin perception

**DOI:** 10.1101/2023.03.15.532805

**Authors:** Monika Kubalová, Karel Müller, Petre Ivanov Dobrev, Annalisa Rizza, Alexander M. Jones, Matyáš Fendrych

## Abstract

The nuclear TIR1/AFB - Aux/IAA auxin pathway plays a crucial role in regulating plant growth and development. Specifically, the IAA17/AXR3 protein participates in root development, and the accumulation of its mutant variant, AXR3-1, which cannot bind auxin, leads to severe root growth phenotype and agravitropism. However, the mechanism by which AXR3 regulates cell elongation is not fully understood. Here we show that the inducible expression of AXR3-1 in the *Arabidopsis thaliana* root triggers excessive cell elongation that is followed by growth arrest of the root. We exploited this effect to reveal the underlying molecular mechanism of AXR3 action. We show that AXR3-1 acts exclusively in the nucleus where it interferes with the nuclear auxin transcriptional pathway, while the rapid cytoplasmic auxin root growth response is not affected. The analysis of the transcriptome of the induced AXR3-1 roots revealed changes in phytohormone perception and homeostasis. We show that the accumulation of AXR3-1 disturbs auxin homeostasis which leads to excessive auxin accumulation. At the same time, the reaction of the AXR3-1 roots to gibberellin is altered. These results show that the IAA17/AXR3 maintains an optimal cell elongation rate by controlling the auxin response, auxin homeostasis and the interplay with gibberellin signaling.

## Introduction

Cell growth is a fundamental part of plant development and responses to external stimuli. The combination of cell division and cell elongation is the dominant factor in plant body formation. In the root tip, cells divide in the meristematic zone (Dolan et al. 1993), and then elongate in the elongation zone, driving the root growth (Beemster and Baskin 1998). Many genetic and environmental factors determine root cell growth, including internal growth regulators – phytohormones (Vanstraelen and Benková 2012).

One of the key hormones that affect root growth is auxin. Auxin is involved in many root developmental processes, from its role in meristem maintenance, through controlling the overall root architecture, to the formation of root hairs and lateral roots (reviewed in Overvoorde et al., 2010). Auxin gradient is associated with patterns of root cell proliferation, elongation, and differentiation. Its concentration is highest around the quiescent center, promoting growth and cell division, and decreases gradually in the shootward direction, leading to cell differentiation (Grieneisen et al. 2007; Mähönen et al. 2014).

Apart from these processes, auxin exerts an inhibitory effect on root cell elongation, which was shown already in 1939 (Bonner and Koepfli 1939). This effect is mediated by modifying cell wall extensibility, membrane depolarization and effects on intracellular signaling processes (Rayle & Cleland, 1992; reviewed in Majda & Robert, 2018). Furthermore, asymmetric redistribution of auxin in the gravistimulated root and its accumulation in the lower part of the root causes inhibition of cell elongation, resulting in bending of the root in the direction of gravity (Friml et al. 2002). How auxin governs such a wide range of responses occurring on long time scales (cell division and differentiation) and on short time scales (cell growth inhibition) is not completely understood. One explanation might be the involvement of the transcriptional and the rapid, non-transcriptional auxin signaling pathways.

It is well established that auxin is perceived in the cell nucleus by the TIR1/AFBs – Aux/IAA coreceptor complex, representing the nuclear auxin pathway. The TIR1/AFB proteins are part of an Skp-Cullin-F-box (SCF) complex that moderates degradation of the Aux/IAA proteins via the 26S proteasome. Upon higher auxin levels, Aux/IAAs-mediated repressor recruitment is hindered, auxin response factors (ARFs) are released, resulting in modulation of transcription (Leyser 2018). In addition, it has been recently suggested that the adenylate cyclase activity of TIR1/AFB is required for the function of the nuclear auxin pathway (Qi et al. 2022).

Alongside the nuclear auxin pathway, auxin triggers changes in plasma membrane potential, cytosolic Ca^2+^ spikes, and a rise in the apoplastic pH. Auxin also activates ROPs (Rho of plants) thereby regulating their downstream effectors such as the cytoskeleton or vesicle trafficking (reviewed in Dubey et al., 2021). In the root, a mutualistic interaction of auxin signaling pathways was also shown. The synergistic functioning of the nuclear auxin pathway and the cytoplasmic AFB1 receptor for fast auxin responses (Dubey et al. 2023; Prigge et al. 2020; Serre et al. 2021) is essential for the regulation of root bending. However, their antagonistic effect is involved in the formation of lateral roots (Dubey et al. 2023). On the contrary, the TMK1-mediated and nuclear auxin pathway act antagonistically in the regulation of root apoplastic pH or root growth inhibition (Li et al. 2021).

In addition to auxin, the role of other plant hormones in root morphogenesis has been widely investigated. For example, gibberellins accumulating in elongating root cells (Rizza et al., 2017; Shani et al., 2013) are required for root cell expansion (Fu and Harberd 2003; Tanimoto 2012; Ubeda-Tomás et al. 2008) and meristem maintenance (Shtin, Dello Ioio, and Del Bianco 2022). The interaction of auxin and gibberellin signaling pathways occurs during several plant developmental processes and at different levels, including mutual effect on biosynthesis (Frigerio et al. 2006; Muir 1962; Ross et al. 2000, 2001; Wolbang and Ross 2001), deactivation (Frigerio et al. 2006), signaling (Frigerio et al. 2006; Li et al. 2015) and transport (Li et al. 2015; Löfke et al. 2013; Willige et al. 2011). The molecular network underlying the auxin-gibberellin crosstalk is still not fully understood.

The semi-dominant *axr3-1* Arabidopsis mutant harbors a stabilizing point mutation in the auxin binding domain II of the IAA17/AXR3 protein. This leads to severe growth and developmental defects, consistent with an auxin resistance (Leyser et al. 1996), including agravitropic and auxin insensitive roots (Knox 2003; Leyser et al. 1996; Li et al. 2009; Lucas et al. 2008; Mähönen et al. 2014; Swarup et al. 2005) and blocking of root hair initiation and elongation (Kim et al. 2006; Knox 2003) or lateral root development (De Smet et al. 2007). The AXR3-1 mutant protein shows a dramatically increased stability while its nuclear localization is not abolished (Ouellet, Overvoorde, and Theologis 2001). In this work, we exploit the mutant AXR3-1 protein as a tool to unravel how Aux/IAAs regulate root growth. By analyzing the line inducible overexpressing AXR3-1, we show that the accumulation of stable AXR3-1 protein leads to a temporary root growth acceleration. To investigate the molecular network and physiological processes occurring in elongating root cells, we use pharmacological and genetical approaches and transcriptomic profiling. Our results provide insights into molecular mechanisms involved in regulation of cell elongation and show the importance of the crosstalk between the gibberellin and auxin signaling pathways.

## Results

### Induction of AXR3-1 causes a promotion of root cell elongation

The severe phenotype of *axr3-1* plants is a result of a long-term accumulation of the AXR3-1 protein, inevitably causing pleotropic secondary effects. To determine the primary effects of a sudden AXR3-1 protein accumulation in the *Arabidopsis thaliana* roots we used an estradiol-inducible *g1090::XVE>>AXR3-1-mCherry* line (*iAXR3-1*; Mähönen et al., 2014). Apart from the described disturbance of auxin sensitivity – root hair growth arrest and agravitropic phenotype (Knox 2003; Leyser et al. 1996; Mähönen et al. 2014) (Fig. **1a,c,d**), AXR3-1 overexpression surprisingly triggered a temporary growth acceleration (Fig. **1b**). Unlike the stable growth rate of the control line, *iAXR3-1* plants displayed accelerated growth over several hours, reaching the maximal growth rate approximately 4-6 h after the start of treatment. Later, the growth rate gradually decreased until root growth stopped (Fig. **1b**). The effect of AXR3-1 induction could be detected after 50 min of estradiol treatment, overcoming the auxin-induced growth inhibition (Fig. **1c**). Interestingly, the growth acceleration preceded the detectable fluorescence of the AXR3-1-mCherry protein, which is likely caused by the time required for the fluorescence protein to mature (Balleza, Kim, and Cluzel 2018).

**Fig. 1.**
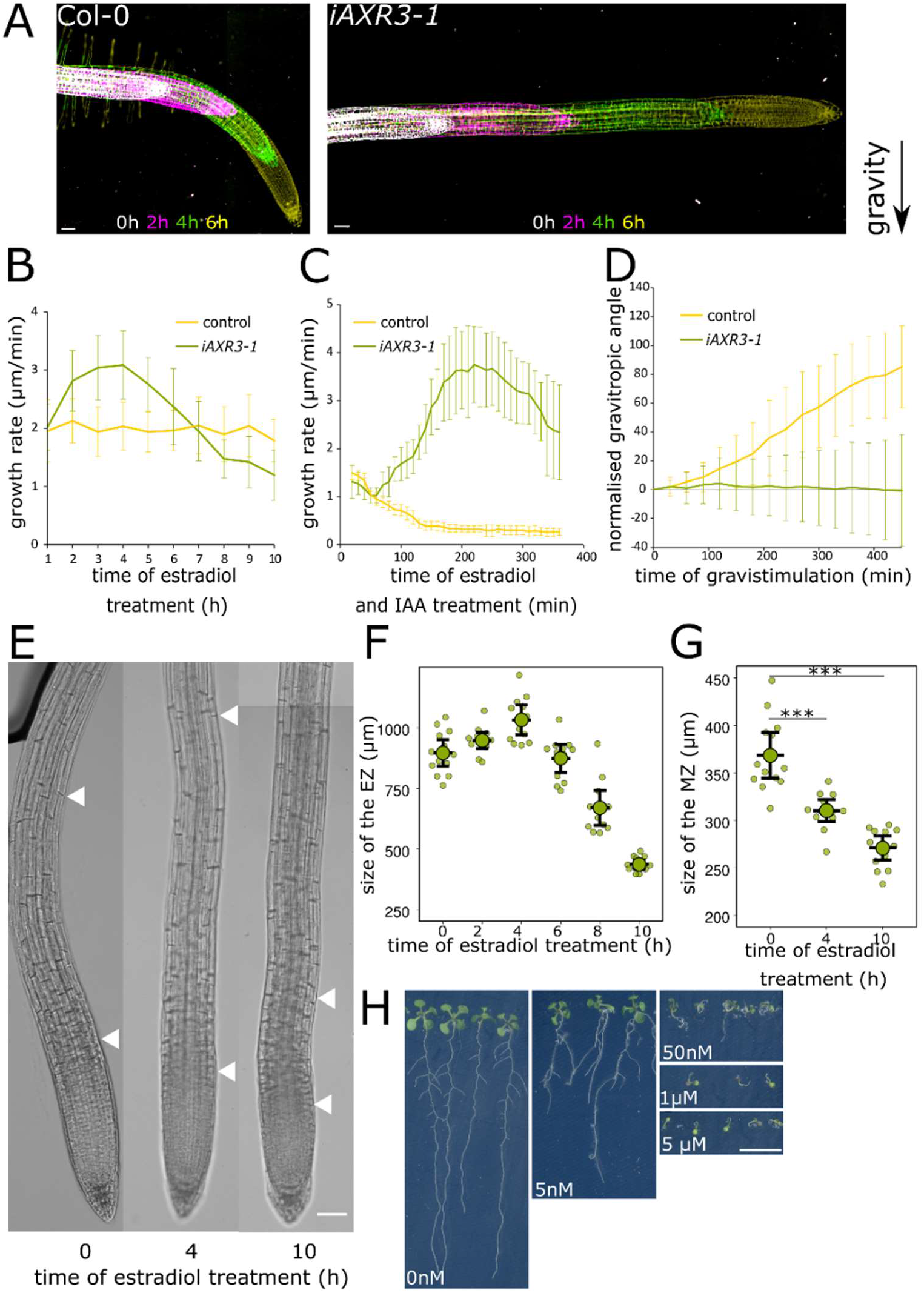
Inducible AXR3-1 overexpression results in increased root cell elongation and altered root morphology. (**a**) Time series of Col-0 and *iAXR3-1* root after 0 h (white), 2 h (magenta), 4 h (green) and 6 h (yellow) of gravistimulation; roots were pretreated with estradiol for 2 h. (**b,c**) Root growth rate of *iAXR3-1* and Col-0 after estradiol treatment (**b**) and 10 nM IAA - estradiol co- treatment (**c**). (**d**) Normalized gravitropic angle of *iAXR3-1* and Col-0 roots after 2 h of estradiol pretreatment. (**e**) Longitudinal zonation of *iAXR3-1* roots after 0, 4 and 10 h of estradiol treatment; arrows indicate the beginning and the end of the elongation zone. (**f-g**) Quantification of the size of the root elongation zone (**f**) and meristematic zone (**g**) of *iAXR3-1* plants after estradiol treatment. (**h**) The long-term effect of increasing estradiol concentration on the morphology of *iAXR3-1* plants; plants grown for 10 d on the indicated concentration of estradiol. Error bars are SD in (**b**), (**c**), (**d**) and CI in (**f**), (**g**); scale bar in (**a**), (**e**) = 50 μm, in (**h**) = 1 cm.

To test how the growth rate changes correlate with longitudinal zonation of the root tip, we determined the meristem and elongation zone size dynamics upon AXR3-1 induction. The meristem size decreased with the increasing time after induction (Fig. **1e,g**). In contrast, the elongation zone size increased, reached the maximum size at 4 h of AXR3-1 induction and then gradually decreased (Fig. **1e,f**). This trend demonstrates that the root growth acceleration can be explained by the elongation zone size changes.

The growth acceleration occurred when *iAXR3-1* plants were treated with at least 1 μM estradiol (Fig. **S1a**), however, *iAXR3-1* roots were agravitropic already at 50 nm estradiol treatment (Fig. **S1b**). A concentration of 5 μM estradiol was used for all subsequent growth promotion experiments to achieve the highest effect on the mutant line with no apparent effect on the control line.

In addition to the short-term effects (hours) of AXR3-1 induction, we analyzed its long-term effect on root growth and morphology. As expected, with the increasing concentrations of estradiol, the accumulation of AXR3-1-mCherry protein increased (Fig. **S1c**). This correlated with a decrease in root length, impaired formation of lateral roots (Fig. **1h**) and root hairs (Fig. **S1d**). Already the low estradiol concentration treatment resulted in shorter, agravitropic roots that lacked lateral roots (Fig. **1h**, **S1b**) indicating different sensitivity of various auxin-related processes to AXR3-1 levels.

When we analyzed the *iaa17/axr3* loss of function mutant, we were unable to detect differences in root growth rate, response to gravistimulation or auxin sensitivity when compared to the Col-0 control line (Fig. **S1e,f,g**). As IAA17/AXR3 is one of 29 Aux/IAA proteins (Reed 2001), this results likely from the redundancy within this protein family.

Taken together, these results suggest the essential role of fine-tuned regulation of auxin signaling in root growth, mediated by IAA17/AXR3 protein. The follow-up question is the subcellular localization of AXR3-1 action.

### Growth acceleration and auxin insensitivity require nuclear localization of AXR3-1

The AFB1 auxin co-receptor and several Aux/IAA proteins localize not only to the nucleus, but also partially to the cytoplasm (Prigge et al. 2020; Zhang et al. 2019), and the AFB1-dependent rapid signaling occurs in the cytoplasm (Dubey et al. 2023; Prigge et al. 2020). We therefore tested whether AXR3-1 might function outside the nucleus and participate in the rapid auxin response. We expressed the wild type IAA17/AXR3 and its stabilized form AXR3-1 under the control of the strong ubiquitous g1090 promoter (Ishige et al. 1999) and the PIN2 promoter, specific for the lateral root cap, epidermis and cortex root cells. Both protein forms localized exclusively to the nucleus (Fig. **2a**), in agreement with Ouellet et al. (2001), and corresponding to the presence of the bipartite nuclear localization signal (NLS) in IAA17/AXR3 sequence (Abel, Oeller, and Theologis 1994). To reveal a potential role of a minor cytoplasmic fraction of IAA17/AXR3, we employed several strategies to modify the localization of the AXR3-1 protein. We attached a nuclear export signal (NESNES, Schwarzerová et al., 2019), or the 8KFarn plasma membrane anchor (Simon et al., 2016) to AXR3-1. However, the AXR3-1 in these lines was still present in the nucleus (Fig. **S2a**). Further, we mutated the bipartite NLS of AXR3-1 using the 31K→E, 32R→S and 207R→G substitutions, creating the AXR3-1ΔNLS (Fig. **2b**; Ouellet et al. (2001)). AXR3-1ΔNLS protein localized to the cytoplasm (Fig. **2c**). In contrast to AXR3-1, the expression of AXR3-1ΔNLS in two independent homozygote lines did not interfere with auxin sensitivity (Fig. **2d**, **S2b**) or gravitropic response (Fig. **2g**), and it did not affect the root phenotype in long-term treatment (Fig. **S2c**). Moreover, the expression of AXR3-1ΔNLS did not lead to growth acceleration. Instead, AXR3-1ΔNLS exerted a mild negative effect on root growth (Fig. **2d**, **S2b**).

**Fig. 2.**
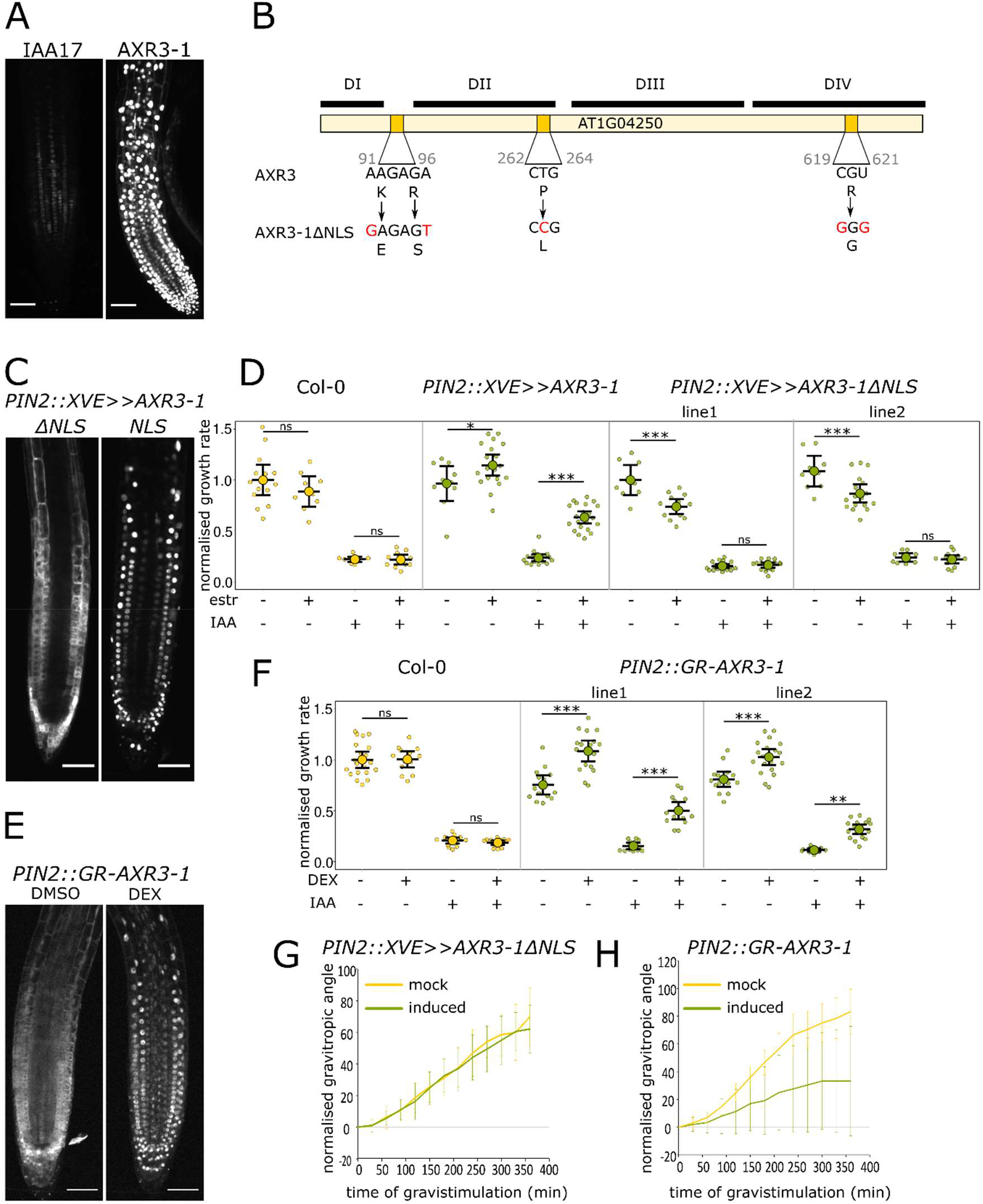
Nuclear localized AXR3-1 is required to affect root growth rate and gravitropic bending. (**a**) Root tip expressing the WT form of IAA17/AXR3 and its stabilized AXR3-1 form driven by the PIN2 promoter. (**b**) Scheme of IAA17/AXR3 coding sequence highlighting domains I - IV and mutations in the auxin-binding DII domain (P>L) and the bipartite NLS (KRR>ESG). (**c,e**) Root tips expressing *PIN2::XVE>>AXR3-1NLS* and *PIN2::XVE>>AXR3-1ΔNLS*, 4 h after estradiol treatment (**c**) or *PIN2::GR-AXR3-1*, 90 min after DMSO/DEX treatment (**e**). (**d, f**) Normalized root growth rate of the indicated lines during a 4 h period in the presence of indicated treatments; (**d**): mock/estradiol and mock/10nM IAA, (**f**): mock/DEX and mock/10 nM IAA. (**g,h**) Normalized gravitropic angle of (**g**) *PIN2::XVE>>AXR3-1ΔNLS*, 2 h of estradiol/DMSO pretreatment and (**h**) *PIN2::GR-AXR3-1*, 2 h of DMSO/DEX pretreatment. Error bars in (**d**), (**f**) are CI and in (**g**), (**h**) are SD; scale bar in (**a**), (**c**), (**e**) = 50 μm.

In addition, we fused the AXR3-1 protein to the glucocorticoid receptor fragment (GR; Schmitt & Stunnenberg, 1993) to allow for a dexamethasone (DEX)-dependent shift in localization between the nucleus and the cytoplasm. Under mock conditions (DMSO), the signal was present in the cytoplasm (Fig. **2e**). Compared to the control, root growth was mildly negatively affected by AXR3-1 localized in cytoplasm (Fig. **2f**). However, after the addition of DEX, the protein moved to the nucleus (Fig. **2e**) and triggered a growth stimulation, auxin insensitivity (Fig. **2f**) and an agravitropic phenotype (Fig. **2h**). In agreement, plants grown on DEX-containing medium for 5 d were agravitropic (Fig. **S2d**). Notably, although the expression of the positive control – the AXR3-1 protein localized in the PIN2 domain led to growth acceleration and auxin insensitivity as expected, the effect on root growth was weaker than in the original *iAXR3-1* plants, where the expression is driven by the g1090 promoter. These results from two independent approaches clearly show that phenotype caused by AXR3-1 overproduction requires AXR3-1 protein localized in the nucleus. This indicates that the observed growth phenotype depends on AXR3-1-mediated transcriptomic changes, rather than its involvement in cytoplasmic rapid responses.

### Induction of AXR3-1 results in upregulation of genes involved in phytohormone and ROS pathways and downregulation of cell wall-related genes

To identify the genes responsible for the growth stimulation after induction of AXR3-1, we analyzed transcriptomic profile from the root tips of two *AXR3-1* inducible lines: estradiol-inducible *iAXR3-1* and the heat shock-inducible *HS::AXR3-1* (Knox 2003). In addition, to identify genes regulated by auxin signaling, we obtained transcriptome of Col-0 root tips treated for 20 min with indole-3-acetic acid (IAA) and auxin signaling inhibitor (PEO-IAA, (Nishimura et al. 2009)). Pairwise comparison identified 906 differentially expressed genes upon AXR3-1 activation, 160 of them are regulated by IAA treatment (Supporting Information - Excel file). In the resulting dataset, we could detect the IAA17/AXR3 gene as one of the most upregulated genes in *iAXR3-1* roots; expression of heat shock proteins was upregulated after heat shock as expected; and the known auxin-inducible genes were induced in the IAA-treated roots (Fig. **S3a**).

To find targets of AXR3-1, we selected candidate genes showing downregulation in *iAXR3-1* and changed expression after IAA treatment. Further, we narrowed the candidate gene list based on their expression and/or described role in the root (using TAIR - Berardini et al., 2015). We obtained a set of 16 genes potentially causing the *iAXR3-1* growth stimulation. To test their involvement in root growth stimulation, we isolated the respective mutants and analyzed the growth phenotypes and additional auxin-related phenotypes. The summary of these experiments and results can be found in Table **S1**. However, none of tested mutants showed contribution to growth promotion, but we observed some auxin dependent phenotypic deviations. MISSE KUSSEI-like mutant showed a shorter root (Fig. **S3b**), the auxin up-regulated F-box protein1 (AUF1) mutant displayed a higher number of lateral roots (Fig. **S3d**) and pectin methylesterase (PME41) mutant showed higher root hair density (Fig. **S3c**).

In accordance with previously published information about ploidy changes in the *axr3-1* mutant plants (Ishida et al. 2010), analysis of the transcriptome data showed a substantially change in genes involved in cell cycle regulation (Fig. **S3e**). Therefore, we compared the ploidy state of root tips of *iAXR3-1* and Col-0 2 h and 20 h after estradiol treatment. The results, however, revealed no difference in ploidy between the control and the *iAXR3-1* line (Fig. **S3f**). This difference could be caused by the different tissue types analyzed (whole plants and root tips).

The fact that the screening of the selected mutants did not reveal a single gene that would control the rapid growth promotion suggests that the regulation is more complex, and it involves groups of genes or processes. Therefore, we performed the gene ontology (GO) analyses (DAVID - Huang et al., 2009). We divided the differentially expressed genes in *iAXR3-1* roots into functional groups based on their molecular and biological function, cell compartment, or the pathway in which they are included (Fig. **3a**).

**Fig. 3.**
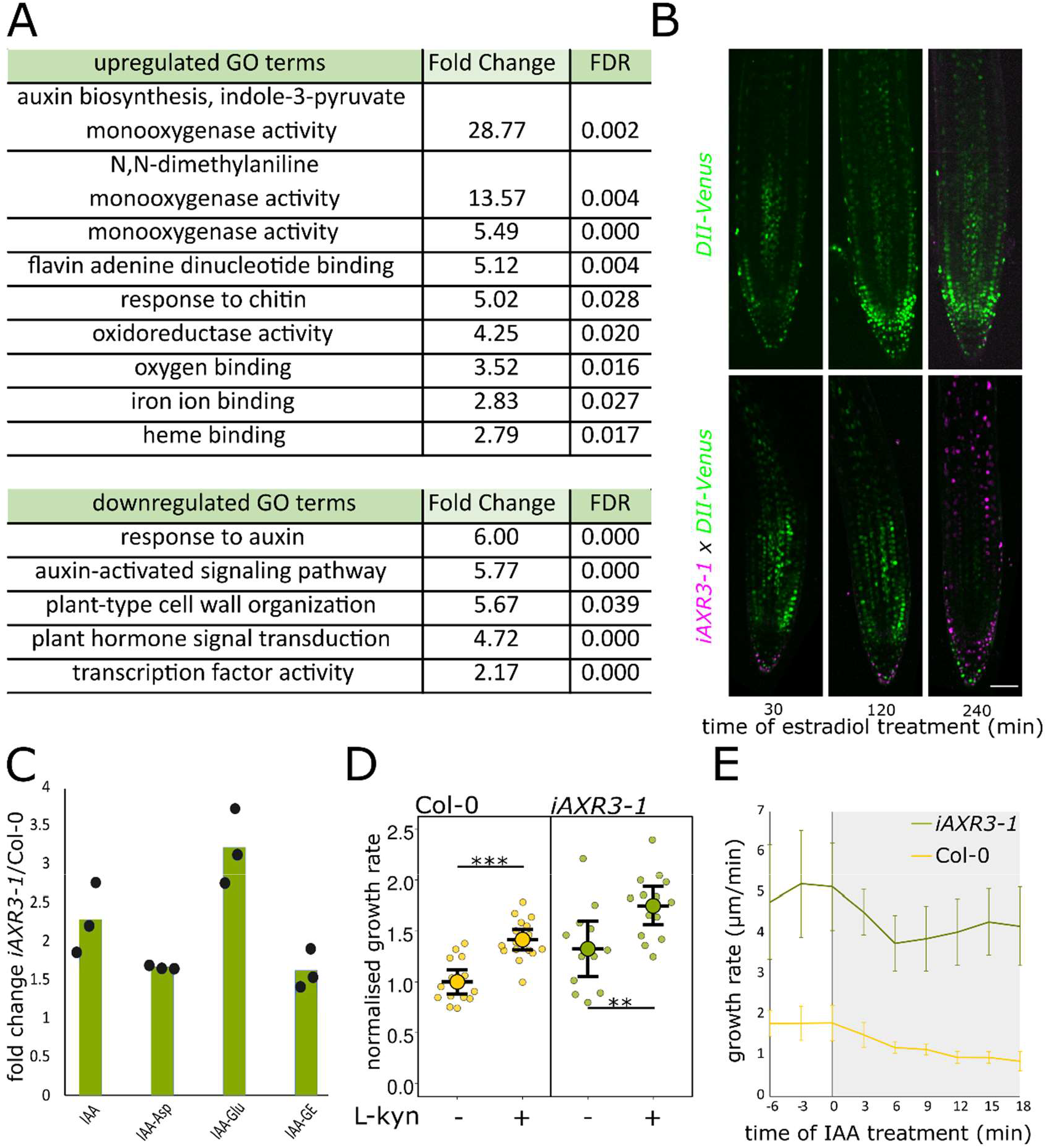
AXR3-1 overexpression affects auxin homeostasis, but auxin sensitivity is partially retained. (**a**) Significantly up/downregulated GO groups upon AXR3-1 induction in root tips, FDR < 0.05. (**b**) Root tips of *iAXR3-1 x DII-Venus* cross after estradiol treatment. (**c**) Fold change of IAA and IAA metabolites enriched in *iAXR3-1* root tips measured by LS-SM hormone assay, IAA = indole-3-acetic acid, IAA-Asp = IAA-aspartate, IAA-Glu = IAA-glutamate, IAA-GE = IAA-glucose ester. (**d**) Normalized growth rate of *iAXR3-1* and Col-0 roots 4 h after L-kyn and estradiol treatment. (**e**) Growth rate of *iAXR3-1* and Col-0 after 10 nM IAA treatment (indicated as grey part), 2 h of estradiol pretreatment. Error bars in (**d**) are CI and in (**e**) are SD; scale bar in (**b**) = 50 μm.

Notably, GO terms related to phytohormones were significantly enriched, including genes related to auxin, gibberellins, brassinosteroids and ethylene (Supporting Information - Excel file). Corresponding to the phenotype of *iAXR3-1* plants, the expression of genes linked to the cell wall modification and gravitropism were changed significantly.

One of the intriguing observations is the significant overrepresentation of genes linked with reactive oxygen species (ROS) (Fig. **3a**), particularly peroxidases. ROS have been shown to be involved in defining the border between meristematic and elongation zone (Tsukagoshi, Busch, and Benfey 2010). Additionally, the mechanism proposed so far (reviewed in Kärkönen & Kuchitsu, 2015) describes the role of ROS in cell wall modifications and consequent changes in the growth response. We therefore analyzed the levels of superoxide and hydrogen peroxide in induced *iAXR3-1* roots using the DHE (dihydroethidium) and BES-H_2_O_2_-Ac staining, respectively. We observed increased levels of both analyzed types of ROS in the rapidly elongating *iAXR3-1* roots in comparison to controls (Fig. **S3g,h**). To test whether ROS production is necessary for the increased *iAXR3-1* cell elongation, we manipulated ROS levels by applying hydrogen peroxide (H_2_O_2_) or the ROS scavenger N-acethyl-1-cysteine (NAC). However, an increase or decrease in ROS levels led to similar root growth responses in both Col-0 and *iAXR3-1* plants (Fig. **S3i,j**). Both treatments triggered a concentration-dependent growth inhibition. The slightly increased sensitivity of the mutant to both treatments at low concentrations can be explained by the increased level of ROS in *iAXR3-1* roots. To test whether increased ROS level could trigger increased growth rate we applied increasing concentrations of H_2_O_2_ to Col-0 roots. Even at very low concentrations, H_2_O_2_ did not lead to increased elongation of the control line (Fig. **S3k**). Together, these data suggest that ROS production on its own cannot steer cell elongation, but the cell elongation might be followed by the increased level of ROS.

### IAA17/AXR3 affects auxin signaling, perception and metabolism and causes auxin overproduction

As the most overrepresented GO categories are related to auxin, we focused on expression changes in auxin-regulated gene groups in *iAXR3-1* roots. In addition to auxin signaling genes, genes involved in auxin biosynthesis were upregulated after AXR3-1 induction (Fig. **3a**). To approximate auxin levels, we crossed *iAXR3-1* plants with the negative auxin reporter *DII-Venus* line (Brunoud et al. 2012). Upon expression of AXR3-1 protein, the DII-Venus signal disappeared (Fig. **3b**), indicating an increased auxin level. In accordance with these results, liquid chromatography–mass spectrometry (LC-MS) measurement of the hormones level in the root tips confirmed a significant increase in the level of IAA and its metabolites in *iAXR3-1* roots (Fig. **3c**).

As it was mentioned, *axr3-1* mutants were identified for their auxin insensitivity (Knox 2003; Leyser et al. 1996). To test whether the expression of AXR3-1 inhibits the rapid non-transcriptional root response to IAA, we used a microfluidic system allowing the change of control to treatment medium in a few seconds (Serre et al. 2021). Similarly to the control, *iAXR3-1* roots responded to addition of auxin by a rapid reduction of growth rate, even though the resulting growth rate was still higher than in the control (Fig. **3e**). Using an alternative method of growth rate analysis after a 20 min IAA treatment, we confirmed that *iAXR3-1* roots retain the rapid auxin response (Fig. **S4a**). Moreover, chemical inhibition of auxin biosynthesis by the auxin biosynthesis inhibitor L-kynurenine (L-kyn, He et al., 2011) led to increased *iAXR3-1* root growth rate (Fig. **3d**), demonstrating that *iAXR3-1* roots maintained the ability to partially respond to endogenous auxin.

In addition to auxin-related genes, AXR3-1 dependent changes in hormone signaling transduction included changed expression of brassinosteroid- and ethylene-related genes. Therefore, we tested the effect of their modified level on the root growth of *iAXR3-1*. Application of bioactive brassinolide (BL) or brassinosteroid synthesis inhibitor brassinazole (BZR) resulted in the same growth response as the control line (Fig. **S4d**), indicating that these hormones are not involved in the rapid growth of the *iAXR3-1* root. However, the *iAXR3-1* mutant was less sensitive to ethylene (Fig. **S4b**) and its precursor (Fig. **S4c**), confirming the interconnection of the auxin and ethylene signaling pathways (Růžička et al. 2007).

The results provide evidence that induction of AXR3-1 protein in the root led to increased auxin biosynthesis and its accumulation in root cells. Although the nuclear auxin signaling pathway is perturbed, the *iAXR3-1* roots did not completely lose the rapid, non-transcriptional auxin signaling. Therefore, the *iAXR3-1* roots can still partially perceive and respond to endogenous and exogenous auxin. Since the interaction of auxin with other hormones has been widely studied (Vanstraelen and Benková 2012), we can assume that such disruption of auxin homeostasis can also lead to modulation of the homeostasis of other hormones.

### Auxin and gibberellin crosstalk in regulation of the root elongation

In accordance with published data (Frigerio et al. 2006), we noticed that IAA treatment and AXR3-1 overexpression led to changes in expression of several gibberellin metabolism genes (Fig. **S4e**). To address the role of gibberellin in the growth stimulation of *iAXR3-1* roots, we biochemically altered gibberellin level in the root. The application of gibberellic acid 3 (GA3) and gibberellic acid 4 (GA4) without previous inhibition of gibberellin biosynthesis had no effect on root growth, regardless of the genotype we studied (Fig. **4c,d**, **S4f**), corresponding to the published literature (Li et al. 2015; Rieu et al. 2008; Tanimoto 1987). Inhibition of gibberellin biosynthesis using two different inhibitors (paclobutrazol - PAZ and uniconazole - Uni (Izumi et al. 1988)) led to a decrease in root growth rate (Fig. **4a,b**) in both control and *iAXR3-1* lines, already after 4 h of treatment. The application of PAZ/Uni from the initial stages of AXR3-1 induction reduced the growth acceleration but did not lead to its complete prevention (Fig. **4a,b**, **S4g**). Interestingly, exogenously added GA3/GA4 rescued the growth rate of PAZ/Uni treated *iAXR3-1* roots. In contrast, GA could not completely recover the growth rate after 4 h in PAZ/Uni treated Col-0 roots (Fig. **4a,b**). From this data, we conclude that *iAXR3-1* plants are hypersensitive to gibberellin. Intriguingly, the expression of gibberellin catabolic and gibberellin synthetic genes was downregulated in *iAXR3-1* (Fig. **S4e**), suggesting altered levels of gibberellin in *iAXR3-1* roots. We therefore introduced the *GPS2 - GIBBERELLIN PERCEPTION SENSOR 2* line into the *iAXR3-1* to estimate gibberellin levels (Alexander Jones, unpublished). We could, however, not detect any difference between gibberellin level in *iAXR3-1* and control roots (Fig. **4e**). Nevertheless, it should be noted that *GPS2* sensor requires a longer time for reversion and therefore cannot detect a reduction in gibberellin level in the short time period that we monitor. Due to technical limitations, we were also unable to detect gibberellin levels using LC-MS method.

**Fig. 4.**
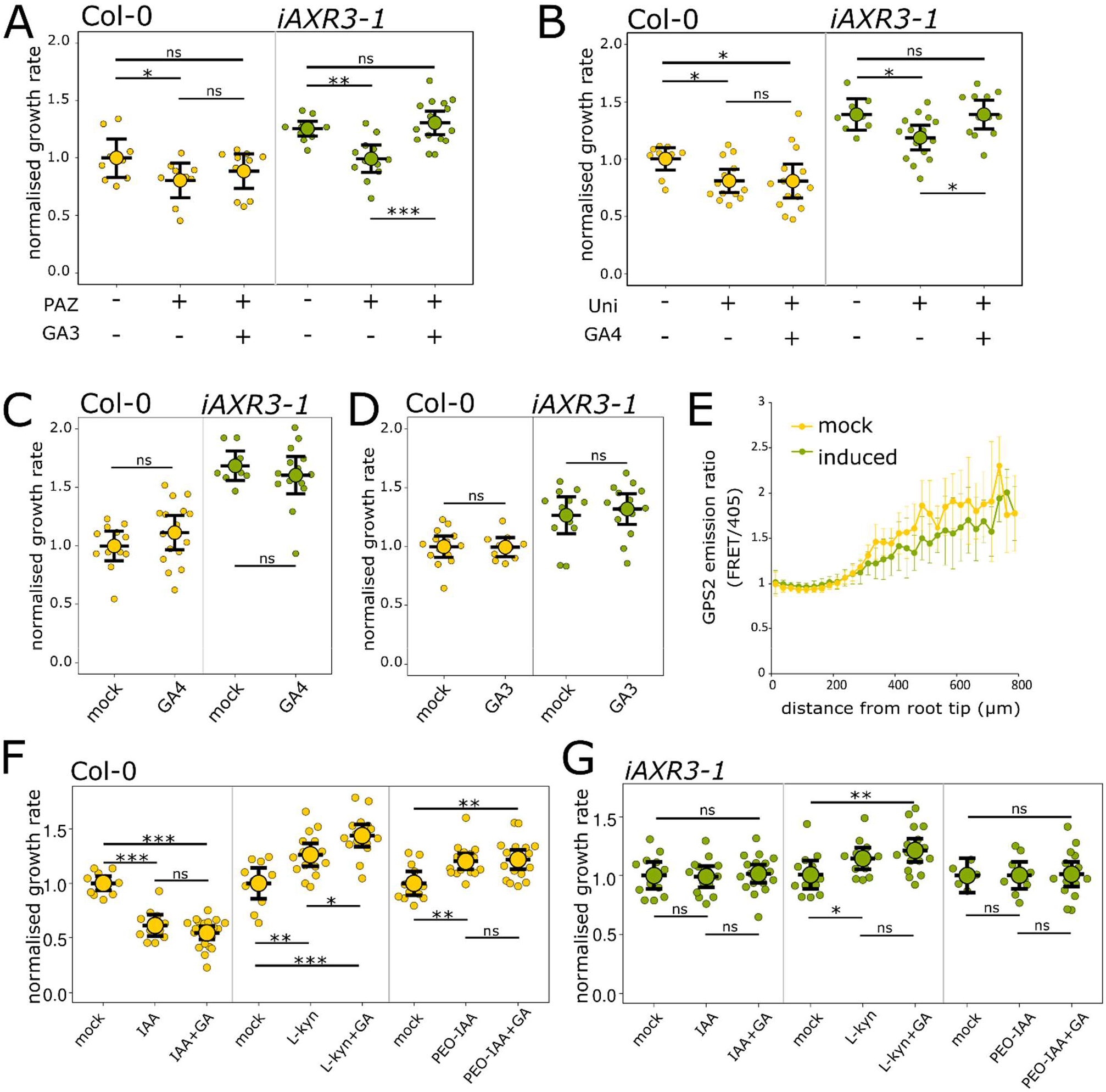
Auxin-gibberellin crosstalk mediated by AXR3-1 regulates root cell elongation. (**a,b**) Normalized growth rate of *iAXR3-1* and Col-0 after 24 h of 1 μM PAZ pretreatment followed by 4 h of 50 μM GA3/1 μM PAZ and estradiol treatment (**a**) and after 6 h of 1 μM Uni pretreatment followed by 4 h of 50 μM GA4/1 μM Uni and estradiol treatment (**b**). (**c,d**) Normalized growth rate of *iAXR3-1* and Col-0 treated by estradiol and 50 μM GA3 (**c**) or 10 μM GA4 for 4 h (**d**). (**e**) Emission ratio of *GPS2 sensor* of *iAXR3-1 x GPS2* roots treated for 4 h with estradiol/DMSO. (**f,g**) Normalized growth rate of Col-0 root (**f**) and *iAXR3-1* roots (**g**) after 4 h of estradiol and 10 nM IAA, 1.5 μM L-kyn, 10 μM PEO-IAA and 50 μM GA3 co-treatment. Error bars in (**a**), (**b**), (**c**), (**d**), (**f**), (**g**) are CI and in (**e**) are SD.

To test the role of the interplay between auxin and gibberellin (Fu and Harberd 2003) in the observed growth stimulation, we treated the plants with bioactive GA3 in combination with pharmacological modification of auxin levels (IAA and L-kyn treatment, respectively), or inhibition of TIR1/AFB pathway (PEO-IAA treatment). In Col-0 plants, the simultaneous application of IAA and GA3 resulted in a non-significant tendency of increased growth inhibition, corresponding with the published data (Li et al. 2015). The comparison of growth rate of PEO-IAA-treated and PEO-IAA+GA3-treated roots displayed no difference. Surprisingly, after the inhibition of auxin biosynthesis (L-kyn treatment) in combination with GA3, there was a more significant promotion of growth (Fig. **4f**), suggesting that auxin prevents the positive effect of gibberellin on root growth. On the contrary, the growth rate of *iAXR3-1* is insensitive to any change caused by the addition of GA3 to IAA, L-kyn or PEO-IAA treated roots (Fig. **4g**).

Although we were not able to determine the gibberellin levels directly, our results show that AXR3-1-dependent modulation of auxin perception and homeostasis increases the root’s sensitivity to gibberellin (Fig. **5**). Taken together, these results suggest an interaction between auxin signaling and gibberellin perception in the root, playing an important role in root cell elongation.

**Fig. 5.**
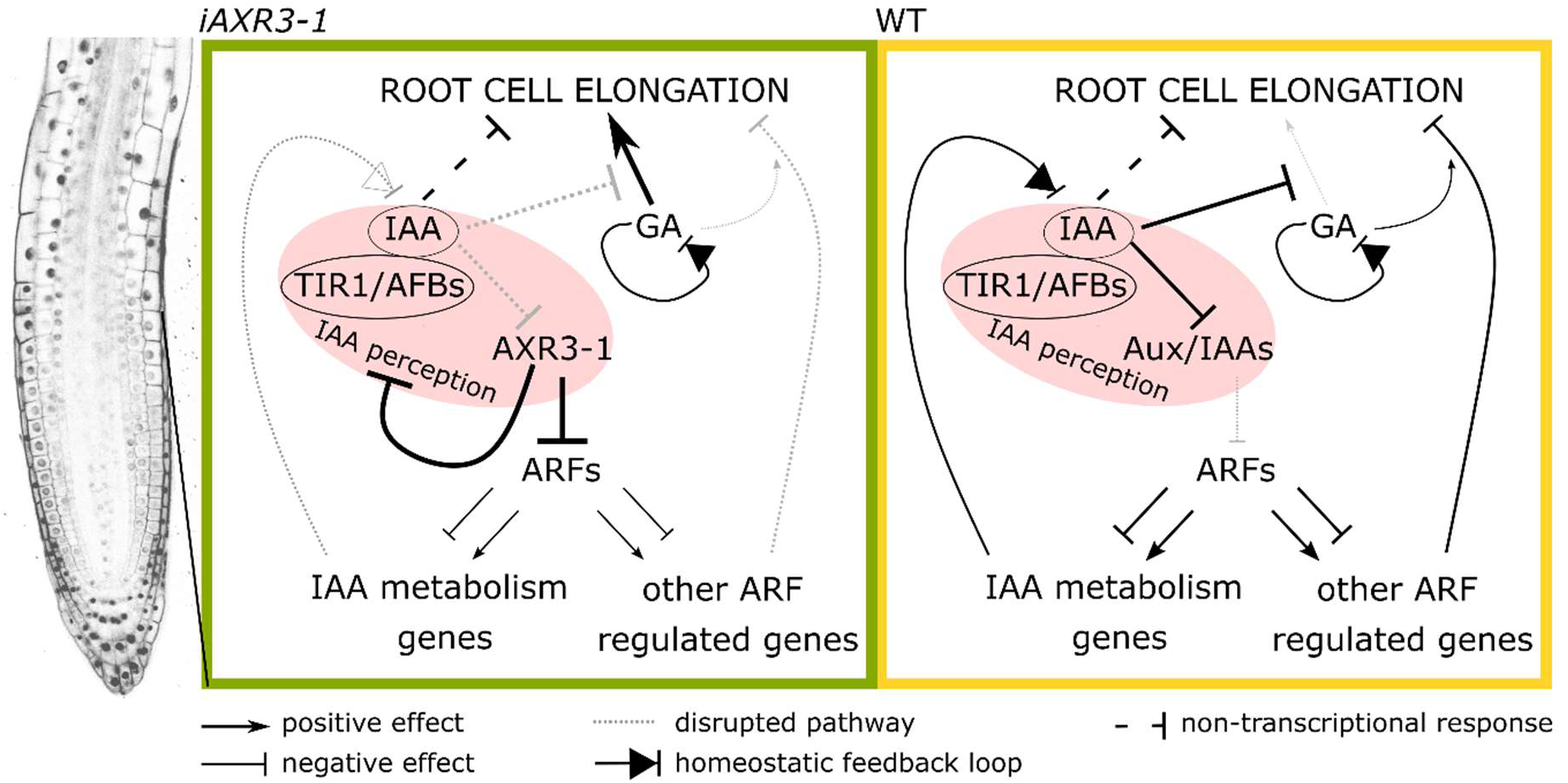
Auxin coreceptor AXR3-1 regulates root cell elongation through modulation of auxin and gibberellin perception. When AXR3-1 is induced, it disrupts auxin perception and alters the modulation of gene expression, including genes involved in root cell elongation. In addition, influencing the expression of IAA metabolic genes leads to disbalance in auxin homeostasis and effect of auxin on cell elongation and on gibberellin perception is reduced. The non-transcriptional auxin effect on cell elongation remains unimpaired. The dysfunction of auxin signaling relieves the inhibitory impact of auxin on gibberellin perception, leading to enhanced responsiveness to gibberellin and contributing to growth promotion. In WT plants, auxin homeostasis is maintained through feedback regulation, whereby auxin regulates the elongation of the cells either by a direct inhibitory effect and at the same time prevents the positive effect of GA on elongation. Pink box highlights the auxin coreceptor complex.

## Discussion

Auxin homeostasis and its perception is essential for optimal plant growth and development. The pool of auxin is maintained by various mechanisms, including regulation of auxin biosynthesis, conjugation, and transport (reviewed in Ljung et al., 2002). Our results demonstrate that proper functioning of the nuclear auxin pathway is essential for regulating root cell elongation and proliferation, thus preventing excessive growth rate. Accumulation of AXR3-1 protein interferes with auxin perception and ARF-modulated gene expression, including the negative transcriptional feedback mechanism controlling the auxin homeostasis (Suzuki et al. 2015). Root cells interpret the AXR3-1 accumulation as low auxin levels, and to establish the proper auxin level, increase the production of auxin. In parallel, the loss of auxin sensing ability allows cells to increase their elongation rate (Fig. **5**). Moreover, as auxin gradient regulates root cell division, expansion, and differentiation (Mähönen et al. 2014; Sabatini et al. 1999), disruption of auxin perception in *iAXR3-1* roots results in disbalance between cell division and elongation, leading eventually to growth arrest. These results indicate that endogenous auxin maintains optimal root growth rate, and that the potential of root cells to elongate is not fulfilled under normal physiological conditions. On the other hand, excessive cell elongation is important to avoid abnormal growth conditions.

Reduction in the overall growth rate of *axr3-1* mutants (Knox 2003; Leyser et al. 1996; Mähönen et al. 2014) is consistent with the observed long-term effect of AXR3-1 protein expression. Hitherto, the impact of short-term AXR3-1 accumulation described here has not been reported, probably because prior studies have primarily examined its effects after 24 h of induction (Mähönen et al. 2014).

Since plants overproducing AXR3-1 are still able to perceive chemical reduction of endogenous auxin level and retain non-transcriptional rapid response to auxin, it is obvious that their capability to perceive auxin is not completely diminished. The differences in sensitivity of individual processes can be explained by the wide range of auxin action, including spatial and temporal aspects and different sensitivity of individual cells to auxin (Sauer, Robert, and Kleine-Vehn 2013). Given that various signaling pathways can exert both cooperative and opposing effects on modulation of root growth and morphology (Dubey et al. 2023; Li et al. 2021), it is plausible to infer that modulation of the nuclear signaling pathway could also affect other auxin cascades. Interestingly, although a rapid non-transcriptional response is required for the early events of gravitropic response (Serre et al. 2021), our data indicate that the cytoplasmic auxin response pathway is not sufficient to promote gravitropic bending in combination with a disrupted nuclear auxin pathway. Furthermore, according to our results, the effect on both root cell elongation and gravitropism is more significant if AXR3-1 is expressed in all root cell types, indicating the contribution of inner cell types to growth control by auxin.

In contrast to the findings obtained using transient expression system (Zhang et al. 2019), we observed solely nuclear localization of AXR3-1. As shown before for the AXR3-1 effect on development (Ouellet et al. 2001), the root growth acceleration and disruption of gravitropism require a nuclear-localized AXR3-1 protein. Interestingly, we observed a weak but consistent effect of cytoplasm-localized AXR3-1 on growth rate. It is intriguing to speculate that the accumulation of the protein in the cytoplasm interferes with the AFB1-dependent auxin response pathway (Dubey et al. 2023; Prigge et al. 2020).

ROS seemed to be one of the possible candidates for the regulation of AXR3-1-dependent growth acceleration. Our results did not prove that hydrogen peroxide application on its own could accelerate root growth. However, it is questionable to what extent pharmacological treatment corresponds to ROS accumulating in different cellular compartments at various concentrations required for the creation of specific signals under physiological conditions (Mhamdi and Van Breusegem 2018). Although we demonstrated an increased level of ROS in *iAXR3-1* roots, this might be associated with a stress response caused by excessive cell elongation (Sharma et al. 2012). Nevertheless, ROS are involved in modulation of cell wall properties (reviewed in Kärkönen & Kuchitsu, 2015). It is a question of future research to investigate AXR3-1-dependent modulation of the cell wall and the possible involvement of ROS.

Increased sensitivity of *iAXR3-1* to GA treatment, the crucial role of gibberellins in regulating root cell expansion (Fu and Harberd 2003; Tanimoto 2012; Ubeda-Tomás et al. 2008), and their accumulation in the elongation zone of the root (Rizza et al., 2017; Shani et al., 2013, this study) makes them a compelling candidates for co-regulators of AXR3-1 growth stimulation. Our results provide strong evidence that gibberellin increases the sensitivity of roots to auxin by acting through AXR3-1 (Fig. **5**). Similarly, GA treatment did not affect growth rate of *tir1* and *axr3-1* mutant (Li et al. 2015) or plants treated with auxin signaling inhibitor (this study).

Due to technical limitations, we were unable to reliably detect any decrease in gibberellin levels within the limited time frame we were focusing on. However, the possibility of AXR3-1-dependent increase in the gibberellin level was ruled out. Thus, there are two possible scenarios: gibberellin level in the roots of *iAXR3-1* is stable or decreased. Despite the fact that there are AXR3-1-dependent changes in the expression of gibberellin metabolic genes, a stable level of gibberellin might be the result of positive feedback loops contributing to gibberellin homeostasis (Yamaguchi 2008). In this scenario, AXR3-1-modulated auxin signaling may increase root cell sensitivity to gibberellin. Second scenario is that increased sensitivity of *iAXR3-1* plants to the level of gibberellin can be caused by decreased gibberellin level in *iAXR3-1* and thus the effect of exogenous application of GA is more pronounced. The reduced level of gibberellin in *iAXR3-1* would, however, be in contrast with data about the crucial role of gibberellin in root development and root cell expansion (Fu and Harberd 2003; Tanimoto 2012). Therefore, the auxin-dependent regulation of the sensitivity of root cells to gibberellin seems more likely to explain *iAXR3-1* phenotype. Regulation of cell sensitivity to gibberellin is of great physiological importance, as shown during germination when lower gibberellin levels trigger root development before the development of the above-ground part begins (Tanimoto and Hirano 2013).

Interestingly, GA stimulates the growth of auxin-depleted plants, revealing the negative effect of auxin on gibberellin perception or response. As AXR3-1 root cells have a decreased auxin perception, the favorable impact of gibberellins on cell elongation is unhindered (Fig. **5**), thus contributing to rapid elongation growth. The fact that AXR3-1-dependent growth stimulation is not completely reduced by inhibition of gibberellin biosynthesis suggests that this phenotype is only partly mediated through the gibberellin response. Another explanation could be that biochemical inhibitors of gibberellin biosynthesis do not completely decrease gibberellin level.

Interference with the nuclear auxin pathway results in disturbance of both auxin homeostasis and the ability to perceive ethylene, gibberellin, and auxin. Our findings offer insights into the connections among distinct auxin signaling pathways and underscore the significance of examining their impacts over varying timeframes. Additionally, we provide evidence that the nuclear auxin signaling pathway regulates the sensitivity of roots to gibberellin, preventing excessive root elongation. These findings contribute to growing evidence of cell expansion regulation by auxin, gibberellin, and their interactions.

## Material and methods

### Experimental material and generation of mutant plants

All lines used are in *Arabidopsis thaliana* Columbia-0 (Col-0) background. We used the following lines: *g1090::XVE>>AXR3-1-mCherry* (Mähönen et al. 2014), *HS::AXR3-1* (Knox 2003), *DII-Venus* (Brunoud et al. 2012), *GPS2* sensor (Alexander Jones, unpublished) and *iaa17* (AT1G04250, SALK_065697C). *iaa17* line was genotyped using following primers: LP CGATTTTCCTCAAGTACGGTG and RP TTTCCTTCACTTGTGCTTTCG. *g1090::XVE>>AXR3-1-mCherry* were crossed with *DII-Venus* and *GPS2* line. Lines *g1090::XVE>>AXR3-1-ΔNLS-mScarlet, PIN2::XVE>>AXR3-1-ΔNLS-mScarlet, PIN2::GR-AXR3-1-mScarlet, PIN2::AXR3-1-mVenusNESNES, PIN2::AXR3-1-mVenus8Kfarn, PIN2::IAA17-mVenus and PIN2::AXR3-1-mScarlet* were prepared in this study as described below.

### Cloning strategy

GoldenBraid methodology (Sarrion-Perdigones et al. 2011) was used as a cloning strategy. For stable transgenic lines, we cloned CDS of IAA17/AXR3 (AT1G04250) or AXR3-1 (88P→L substitution) driven by the PIN2 promoter (1.4 kb upstream of the AT5G57090) and fused with mVenus (*PIN2::IAA17-mVenus*), mScarlet-I (Bindels et al., 2017, *PIN2::AXR3-1-mScarlet*), mVenusNESNES (Schwarzerová et al. 2019) or mVenus8Kfarn (Simon et al. 2016) to C terminus (*PIN2::AXR3-1-mVenusNESNES* and *PIN2::AXR3-1-mVenus8Kfarn*). These constructs were terminated by 35S terminator and cloned into alpha1 vector.

For estradiol-inducible lines, XVE (Zuo, Niu, and Chua 2000) was cloned under the control of the PIN2 promoter (1.4 kb upstream of the AT5G57090) or g1090 promoter (Zuo et al. 2000) terminated by the RuBisCo terminator from *Pisum sativum* and cloned into alpha 1-1 vector. We cloned CDS of IAA17/AXR3, AXR3-1 or AXR3-1-ΔNLS (31K→E, 32R→S and 207R→G substitutions) downstream of the 4xLexA Operon driven by CaMV 35S minimal promoter (Sarrion-Perdigones et al. 2013) and terminated by the 35S terminator, fused with mScarlet-I (Bindels et al. 2017) to the C-terminal part and terminated by 35S terminator. For constructs containing glucocorticoid receptor (*PIN2::GR-AXR3-1-mScarlet*), GR was fused to N-terminus of AXR3-1. These constructs were cloned into the alpha1-3 vectors (Dusek et al. 2020). The alpha transcriptional units were then interspaced with matrix attachment regions (Dusek et al. 2020). Alpha vectors were combined with a Basta resistance cassette and cloned into the pDGB3omega1 binary vector (Sarrion-Perdigones et al. 2013).

All constructs were transformed into the Col-0 ecotype using the floral dip method (Clough & Bent, 1998). All used primers and sequences are listed in Table **S2**.

### Growth conditions and treatments

Seeds were surface-sterilized with chlorine gas (Lindsey III et al. 2017), and stratified for 2 days at 4°C. Seedlings were growth vertically on plates containing 1 % (w/v) agar (Duchefa) with ½ Murashige and Skoog (MS, Duchefa, 0,5g/l MES, 1 % (w/v) sucrose, pH 5.8 adjusted with 1M KOH). Growth chamber conditions were 23°C by day (16 h), 18°C by night (8 h), light intensity of 120 μmol photons m^-2^ s^-1^, 60 % humidity.

The chemicals used to prepare treatments are given in Table **S3**.

To treat the plants, 4-5 d old plants were transferred to treatment medium. Used concentrations and exact treatment times for specific experiments are given in the legend of each figure. In some cases, seeds were germinated on medium containing DEX or estradiol. Control medium always contained the respective mock treatment.

### Microscopic imaging

For high-resolution imaging (including root growth rate measurement over time), vertical stage (von Wangenheim et al. 2017) Zeiss Axio Observer 7 coupled to a Yokogawa CSU-W1-T2 spinning disk unit with 50 μM pinholes and equipped with a VSHOM1000 excitation light homogenizer (Visitron Systems) was used. Images were acquired using the VisiView software (Visitron Systems, v4.4.0.14). The roots were imaged with a Plan-Apochromat 10x/0.45 M27 and Plan-Apochromat 20x/0.8 M27 objectives. We used 515 nm laser for mVenus (excitation 515, emission 520-570 nm), 561 nm laser for mScarlet, mCherry and PI-stained samples (excitation 561, emission 582-636 nm) and 488 nm laser for DHE-/ BES-H_2_O_2_-Ac-stained samples (excitation 488, emission 500-550 nm). For *GPS2* sensor, sample were imaged by Zeiss LSM 880, C-Apochromat 40x/1.2 Imm Corr DIC objective using 515 nm and 405 nm laser.

Low-resolution imaging (gravitropic analysis, root growth rate measurement after 4 h) was performed by vertically placed flatbed scanner (Perfection V700, Epson) with the Epson Scanner software v3.9.2.1US.

### Image analyses and measurements

Software ImageJ Fiji (Schindelin et al. 2012) was used for all image analysis.

For measuring **root growth rate**, distance between root tip positions in consecutive frames was calculated. FiJi plugin Correct 3D drift was used to stabilize the drift of the root tip.

To measure **gravitropic bending**, seedlings were transferred to plates containing the desired media. After 30 min for recovery (or 2 h for treatment), plates were turned 90° and imaged every 30 min. Analysis was done by ACORBA software (Serre and Fendrych 2022). The **size of the meristematic** and **elongation zone** was measured on roots stained with 2,5 μM PI for 15 min. The elongation zone was measured on a time series, whereby the end of the elongation zone was set as the last growing cell. The size of the meristematic zone was measured as a distance from the root tip to the last isodiametric cell.

The **intensity of the fluorescent proteins** was measured either as the intensity of the entire area from the root tip including the elongation zone in the case of mCherry or as the intensity of individual nuclei after removing the background intensity as in the case of the *GPS2* sensor. To analyse **the level of gibberellin**, F1 cross *GPS2 x g1090::XVE>>AXR3-1-mCherry* were induced for 4 h. Gibberellin-dependent ratio change was calculated as FRET/405 fluorescence intensity.

Dihydroethidium (DHE, 5 mM stock in DMSO, Fisher scientific) and BES-H_2_O_2_-Ac (1 mM stock in DMSO, Wako) were used to **stain ROS**. After incubation in dark for 30 min, the signal intensity of epidermal cells was measured.

**Microfluidic experiment** for rapid growth inhibition was done as described in Serre et al. (2021).

### Transcriptomic and gene expression analysis

To obtain transcriptomic data, RNA from 50 root tips (1-2 mm) of *g1090::XVE>>AXR3-1-mCherry* and *HS::AXR3-1* were extracted following the protocol (Plant Total RNA Mini Kit, QIAGEN). *g1090::XVE>>AXR3-1-mCherry* were transferred for 2 h on estradiol-containing medium and then harvested. *HS::AXR3-1* plants were treated for 40 min with 37 °C and harvested after another 80 min of recovery in cultivation room. Each of these lines had an appropriate control (treated in the same way as the mutant line). Col-0 was transferred to medium containing 10 nm IAA or 10 μM PEO-IAA or mock (ethanol) for 20 min. mRNA was prepared from total RNA, followed by Eukaryotic Strand-Specific Transcriptome Library preparation. RNA was sequenced by Illumina PE150 sequencing.

Quality of reads processed by the sequencing service provider was assessed by FastQC software. Transcript abundances (transcripts per million – TPM) were determined using Salmon v1.3.0 (Patro et al. 2017) with parameters --validateMappings, --seqBias, --gcBias, --posBias, --numBootstraps 30. Reference index was built from *Arabidopsis thaliana*, TAIR10 cds library, version 20101214. Statistical evaluation and quality control of data analysis was done using Excel. Transcripts with the following parameters: standard deviation TPM of all replicas/avg of all replicas ≤ 0.6 and log_2_ fold-change ≥1 (upregulated) or ≤ -1 (downregulated) were considered to be significantly differentially expressed. Gene ontology analysis was done using DAVID bioinformatics resources (Huang et al. 2009). GOs with FDR ≤ 0.05 and fold-change ≥ 2 (upregulated) or ≤ -2 (downregulated) were considered to be significantly differentially expressed.

### Selection of candidate genes and the phenotyping of mutants

Candidate genes for further analysis were selected based on the degree of change in their expression upon AXR3-1 overexpression and/or after IAA and PEO-IAA treatment. Genes showing root-specific expression were selected. Mutants were ordered from NASC and subsequently verified by genotyping. The list of individual mutants including the primers used is in Table **S4**.

For the **ploidy analysis**, we followed a protocol from Urfus et al. (2021).

### Liquid chromatography – mass spectrometry measurement

The endogenous phytohormones in root tips were determined according to Prerostova et al., 2021. In brief, about 100 root tips were collected into 100 μl acetonitrile/water (1/1, v/v), spiked with stable isotope labeled internal standards (1 pmol/sample) and were homogenized with 1.5 mM zirconium beads using a FastPrep-24 homogenizer (MP Biomedicals, CA, USA), and incubated at 4°C for 30 min. After centrifugation at 30000x g for 20 min, the supernatant was concentrated to half of initial volume in vacuum concentrator and applied to SPE Oasis HLB 10mg 96-well plate (Waters, Milford, MA, USA). The SPE 96-well plate was washed three times with 100 μl water, followed by elution with 100 μl 50% acetonitrile/water. Aliquot of 5 μl of the eluate was injected into the LCMS system consisting of UHPLC 1290 Infinity II (Agilent, Santa Clara, CA, USA) coupled to 6495 Triple Quadrupole mass spectrometer (Agilent). MS analysis was done in MRM mode, using isotope dilution method. Data acquisition and processing was done with Mass Hunter software B.08 (Agilent).

### Statistical and graphic analyses

Each experiment was performed in a minimum of three biological replicates. Line graphs correspond to a minimum of 7 roots. To compare multiple samples, we used One-Way ANOVA followed by Tukey HSD for normally distributed data. If data did not follow normal distribution, we used Kruskal Wallis test followed by post-hoc Dunn’s test. To compare two sets of normally distributed samples we used Student test. P-value 0.05 > ns, ≤ 0.05 *, ≤ 0.01 **, ≤ 0.001 ***.

All data points in boxplots are shown as dots, the largest dot represents mean. Line graphs and data analysis was done in Microsoft Excel, boxplots were made by SuperPlotsOfData (Lord et al. 2020). Figures were assembled in Inkscape.

## Data availability

Data used to generate figures are attached as Source data file.

## Author contribution

MF and MK conceived and designed the experiments and wrote the manuscript. MK performed the experiments. KM analyzed transcriptomic data. PID performed LC-MS assay. AR and AJ provided GPS2 sensor.

## Acknowledgments

M.F. team received support from the European Research Council (Grant No. 803048). M.K. was supported by Charles University Grant Agency (Grant No. 337021). The authors are grateful to Jan Petrášek, Matouš Glanc for critical reading the manuscript and for discussion and Eva Medvecká for lab support. Confocal laser scanning microscopy performed in the Microscopy Core Facility of Faculty of Science was co-financed by the Czech-BioImaging large RI project LM2023050.

## SUPPLEMENTARY MATERIAL

**Fig. S1.**
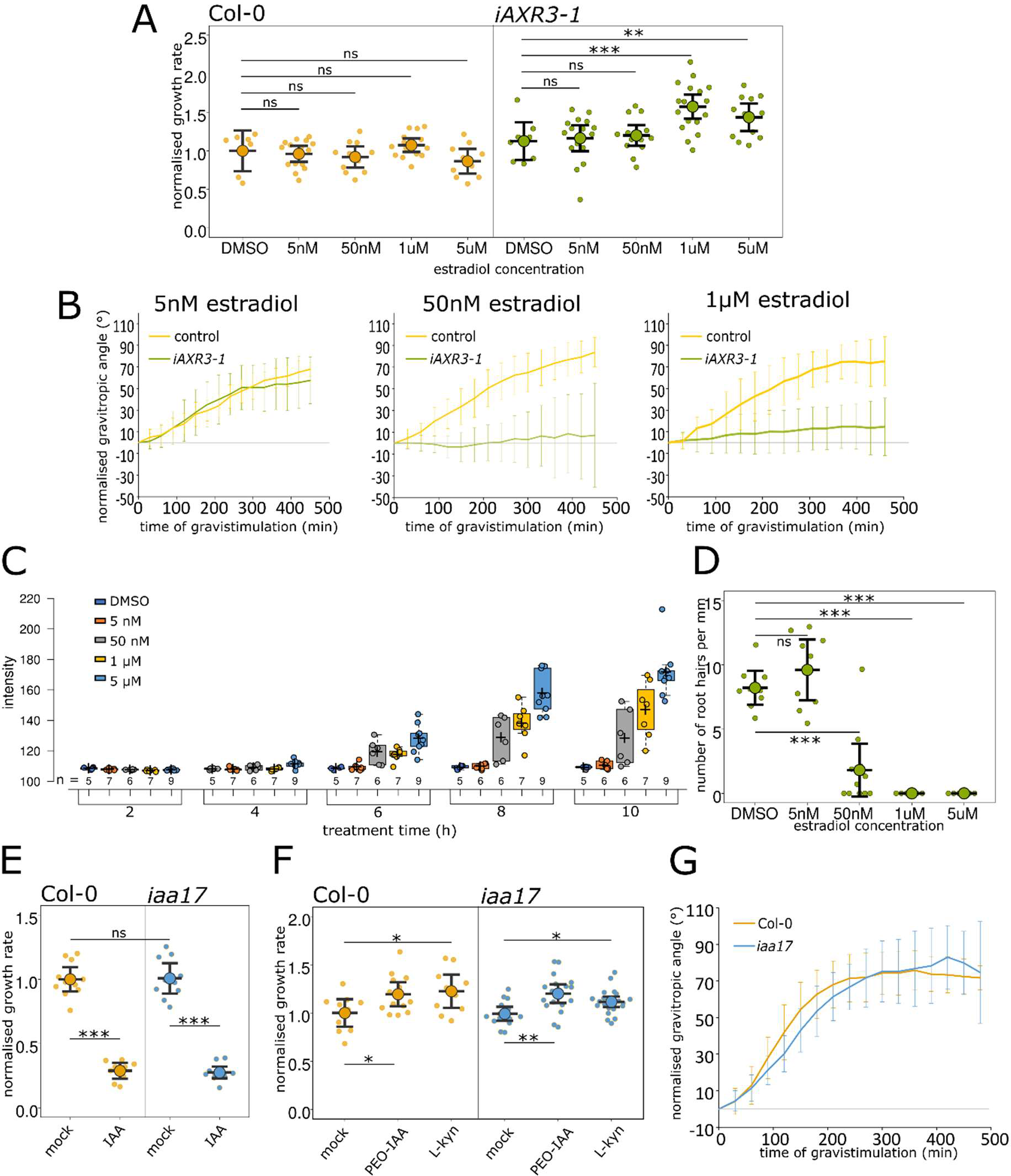
(**a**) Normalized root growth rate of Col-0 and *iAXR3-1* roots treated with increasing concentration of estradiol for 4 h. (**b**) Normalized gravitropic angle of *iAXR3-1* roots treated by the indicated concentration of estradiol, 2 h of estradiol pretreatment. (**c**) Intensity of AXR3-1-mCherry fluorescence in the root tips after induction for the indicated time by the indicated concentrations of estradiol. (**d**) Root hair density (nr. of hairs per mm) of 5 d old *iAXR3-1* roots after induction by different concentrations of estradiol. (**e,f**) Normalized growth rate of *iaa17* mutant after 4 h of 10 nM IAA (**e**) and 1.5 μM L-kyn and 10 μM PEO-IAA treatment (**f**). (**g**) Normalized gravitropic response of *iaa17* and Col-0. Error bars in (**a**), (**d**), (**e**), (**f**) are CI and in (**b**), (**g**) are SD.

**Fig. S2.**
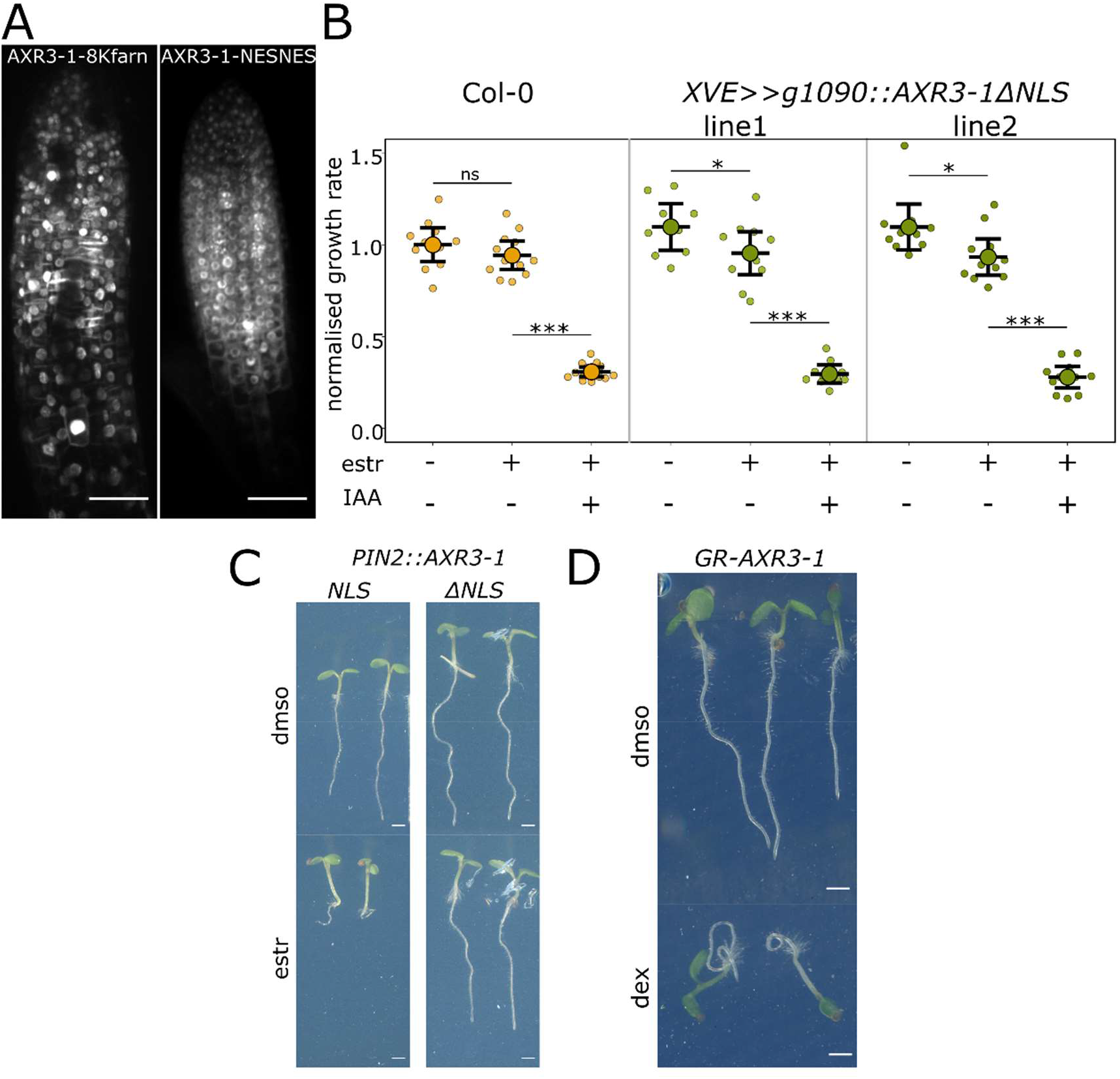
(**a**) Root tip expressing AXR3-1 protein fused with NESNES and plasma membrane anchor 8Kfarn. (**b**) Normalized growth rate of *g1090::XVE>>AXR3-1ΔNLS*, during 4 h of estradiol and 10 nM IAA treatment. (**c**) 5 d old *PIN2::XVE>>AXR3-1NLS* and *PIN2::XVE>>AXR3-1ΔNLS* plants grown on mock/estradiol-containing medium. (**d**) 5 d old *PIN2::GR-AXR3-1* plants grown on DMSO/DEX-containing medium. Error bars in (**b**) are CI, scale bar in (**a**) = 50 μm and in (**c**), (**d**) = 1 mm.

**Fig. S3.**
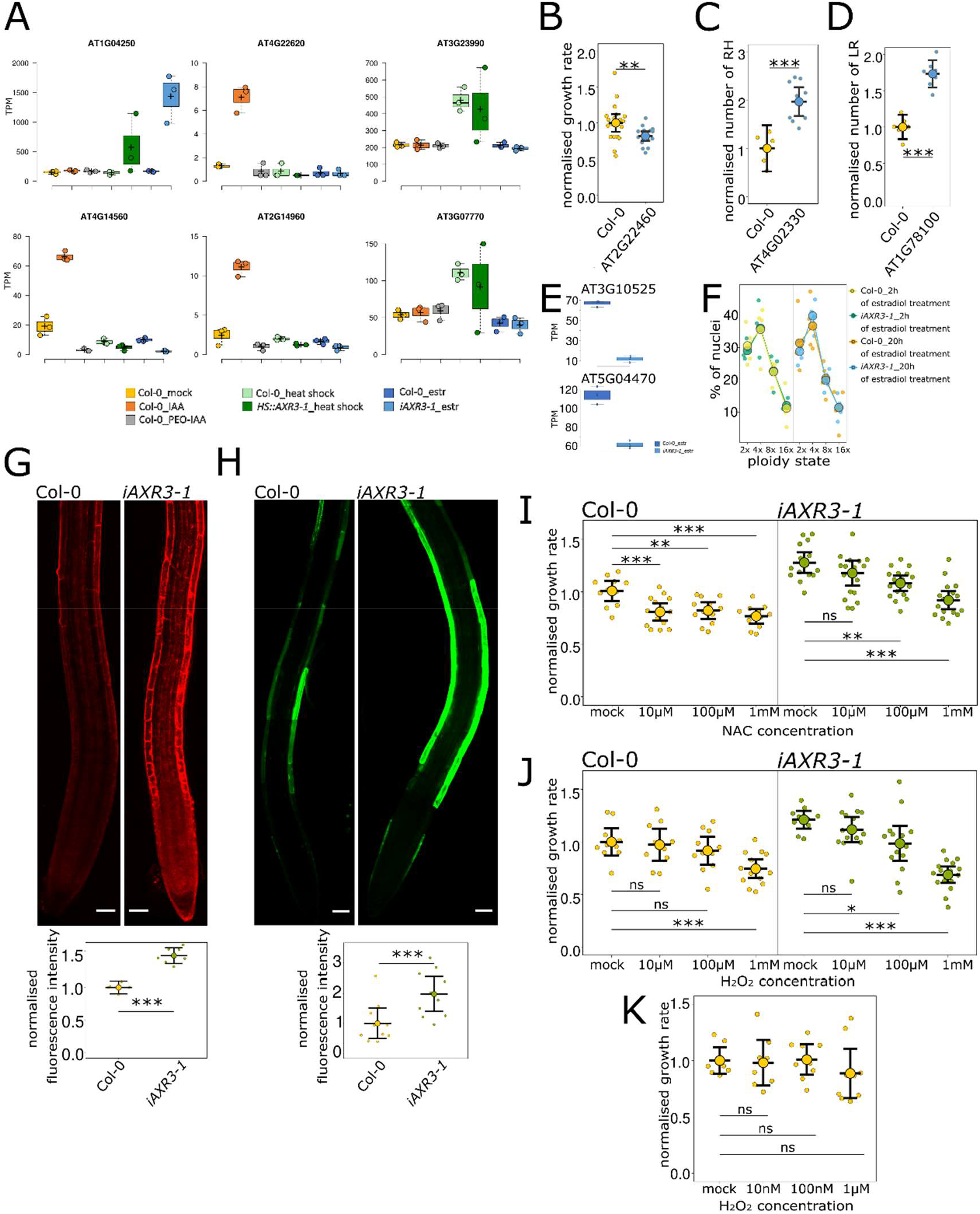
(**a**) Transcripts per millions (TPM)/expression level of genes AT1G04250 (IAA17/AXR3), AT2G14960 (GH3.1), AT4G22620 (SAUR34), AT4G14560 (IAA1/AXR5), AT3G23990 (HSP60) and AT3G07770 (HSP89.1) in Col-0, *iAXR3-1* or *HS::AXR3-1*, experimental details are described in the methods. (**b**) Normalized growth rate of AT2G22460 (MIZU-KUSSEI-like protein) after 4 h growing on MS medium. (**c**) Root hair density of AT4G02330 (PME41) and (**d**) lateral roots of AT1G78100 (AUXIN UP-REGULATED F-BOX PROTEIN 1). (**e**) TPM of AT3G10525 (SMR1) and AT5G04470 (SIM) genes in Col-0 and *iAXR3-1*. (**f**) Ploidy state of root tip cells of Col-0 and *iAXR3-1* 2 h and 20 h after estradiol treatment. (**g,h**) Root tips stained with ROS sensitive dyes DHE (superoxide staining) (**g**) and BES-H_2_O_2_-Ac (hydrogen peroxide staining) (**h**), and the quantification of fluorescence intensity in epidermal cells. Root tips were treated 4 h with estradiol and stained for 30 min. (**i,j**) Normalized growth rate of Col-0 and *iAXR3-1* roots treated with estradiol and different concentration of ROS scavenger NAC (**i**) or H_2_O_2_ (**j**) for 4 h. (**k**) Growth rate of Col-0 roots treated with different H_2_O_2_ concentration for 90 min. Error bars in (**b**), (**c**), (**d**), (**g**), (**h**), (**i**), (**j**), (**k**) are CI, scale bar in (**g**), (**h**) = 50 μm.

**Fig. S4.**
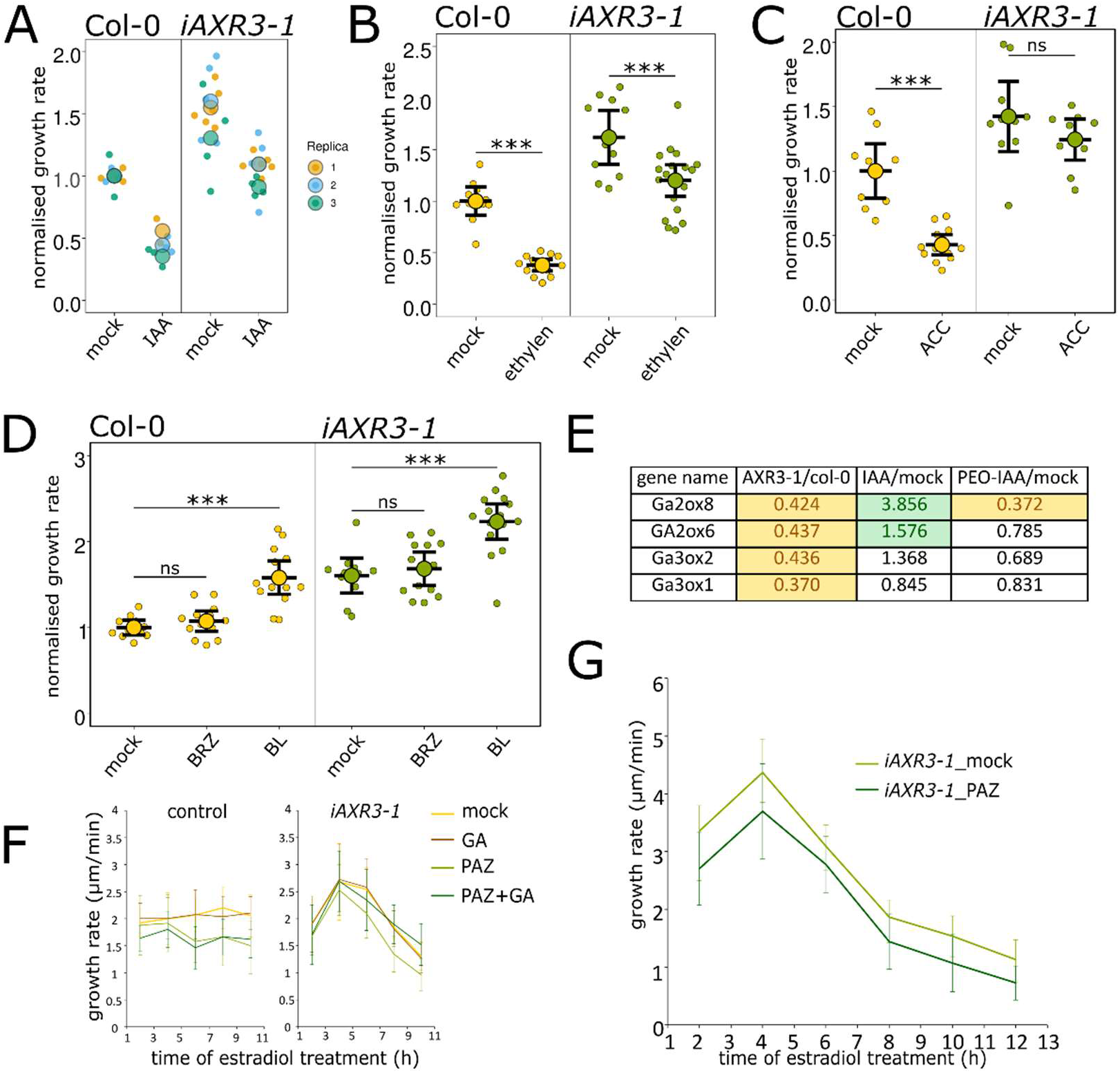
(**a-d**) Normalized growth rate of *iAXR3-1* and Col-0 roots pretreated with estradiol for 2 h and treated for 20 min with 10 nM IAA (**a**), for 4 h with 150 ppm ethylene (**b**), 10 μM ACC (**c**) or 250pM BL and 1μM BZR (**d**). (**e**) Fold change of gibberellin-metabolism genes regulated upon AXR3-1 induction or IAA/PEO-IAA treatment. (**f**) Growth rate of Col-0 and *iAXR3-1* after estradiol, 1 μM PAZ and 50 μM GA3 treatment, without pretreatment. (**g**) Growth rate of *iAXR3-1* roots treated with estradiol and 1 μM PAZ, pretreated with 1 μM PAZ/mock for 24 h. Error bars in (**b**), (**c**), (**d**) are CI and in (**f**), (**g**) are SD.

## SUPPLEMENTAL TABLES

**Table S1.** List of analyzed candidates including their enrichment in *iAXR3-1* or in IAA-treated Col-0 roots and the list of experiments.

| AT code | Name | Fold change |  | Analysis |
| --- | --- | --- | --- | --- |
|  |  | *iAXR3-1*/ con | IAA/mock |  |
| AT5G66580 | PADRE protein | 0.30 | 2.28 | • Root growth rate (see Fig. S3b)<br>• 10 nM IAA/ 10 μM PEO-IAA/ 1.5 μM L-kyn treatment<br>• Number of lateral roots*(see Fig S3d)<br>• Number of root hairs*(see Fig S3c)<br>• Trichome and veins patterning*<br>• Length of etiolated hypocotyl*<br>• Gravitropic response |
| AT5G52750 | Heavy metal transport/detoxification superfamily protein | 3.46 | 2.84 |  |
| AT5G19790 | Related to AP2.11 RAP2.11 | 0.18 | 1.05 |  |
| AT5G12940 | Leucine-rich repeat (LRR) family protein | 0.45 | 1.51 |  |
| AT5G09440 | Exordium like 4 | 0.14 | 1.37 |  |
| AT5G04970 | Pectin methyl esterase inhibitor | 0.44 | 1.00 |  |
| AT4G35210 | hypothetical protein | 0.12 | 2.14 |  |
| AT4G35200 | hypothetical protein | 0.16 | 1.55 |  |
| AT4G02330 | Pectin methyl esterase | 8.37 | 1.86 |  |
| AT3G51410 | hypothetical protein | 0.38 | 2.07 |  |
| AT3G50900 | hypothetical protein | 0.37 | 1.33 |  |
| AT3G25930 | Adenine nucleotide alpha hydrolases-like superfamily protein | 0.14 | 1.16 |  |
| AT2G22460 | MIZU-KUSSEI-like protein | 0.12 | 1.65 |  |
| AT1G80240 | DUF-642 l-gall responsive gene 1 | 0.18 | 1.86 |  |
| AT1G78100 | AUXIN UP-REGULATED F-BOX PROTEIN1 | 0.29 | 3.09 |  |
| AT1G64795 | hypothetical protein | 337.10 | 1.00 |  |

\*The number of lateral roots of 10 d old plants and the number of root hairs of 5 d old plants (grown on MS medium) were manually counted. The length of the etiolated hypocotyl was measured on plants whose seeds were exposed to light for 2 h and then grown vertically in the dark for 3 d. Patterning of trichomes was monitored with a binocular microscope (Stemi 305 EDU Microscope Set/ Olympus SZX7) on 10 d old seedlings, and patterning of veins was monitored on 4 h bleached cotyledons (with 90 % acetone, -20°C) of 10 d old plants.

**Table S2.**
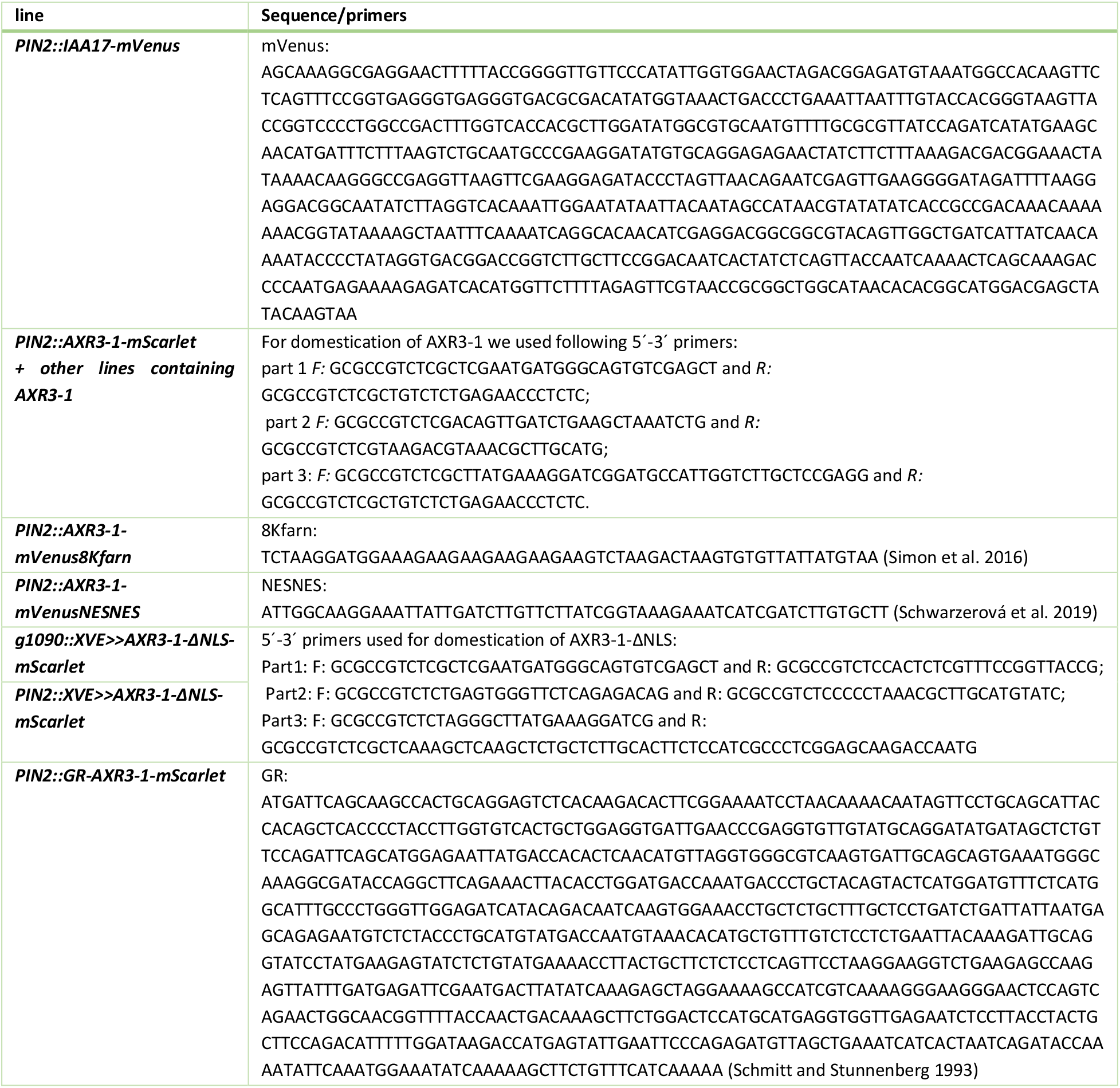
List of primers and sequences used for cloning.

**Table S3.**
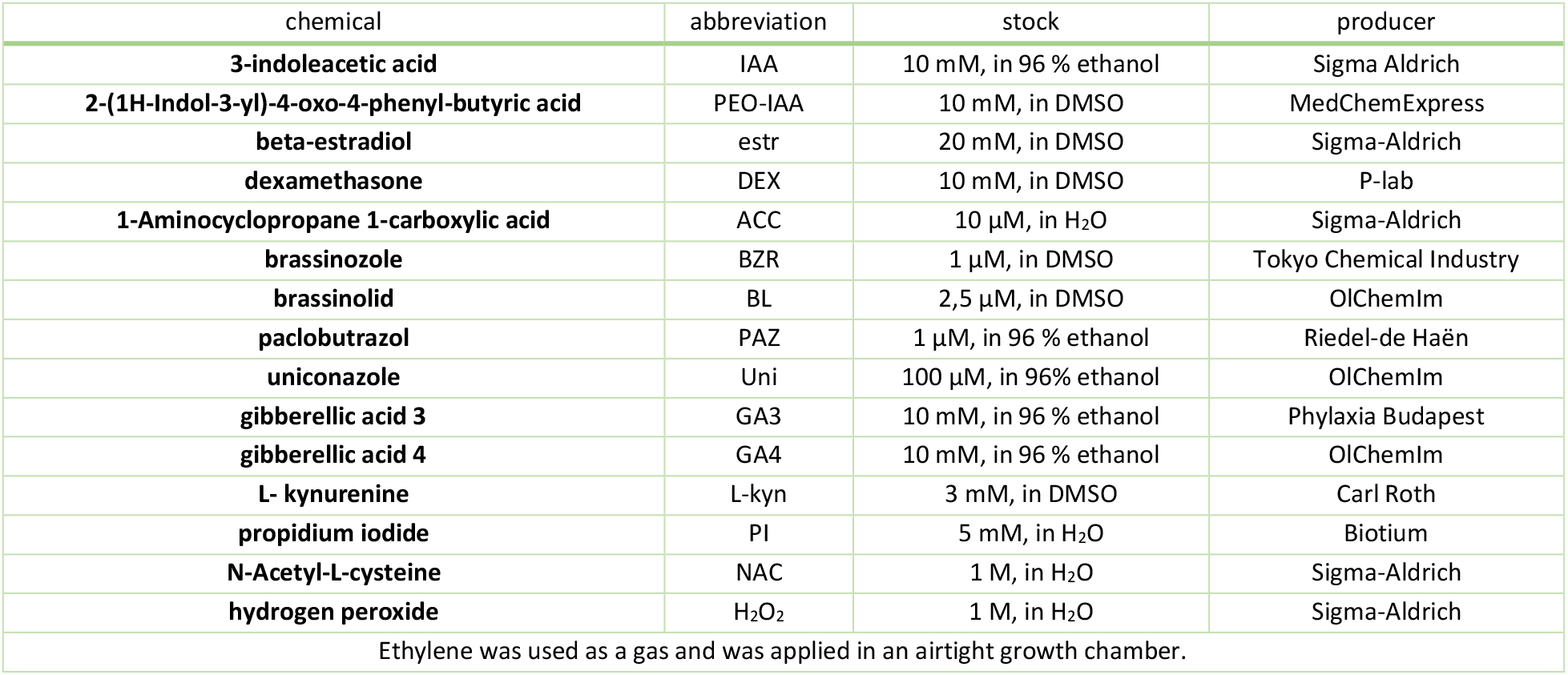
List of chemicals used for treatments.

**Table S4.**
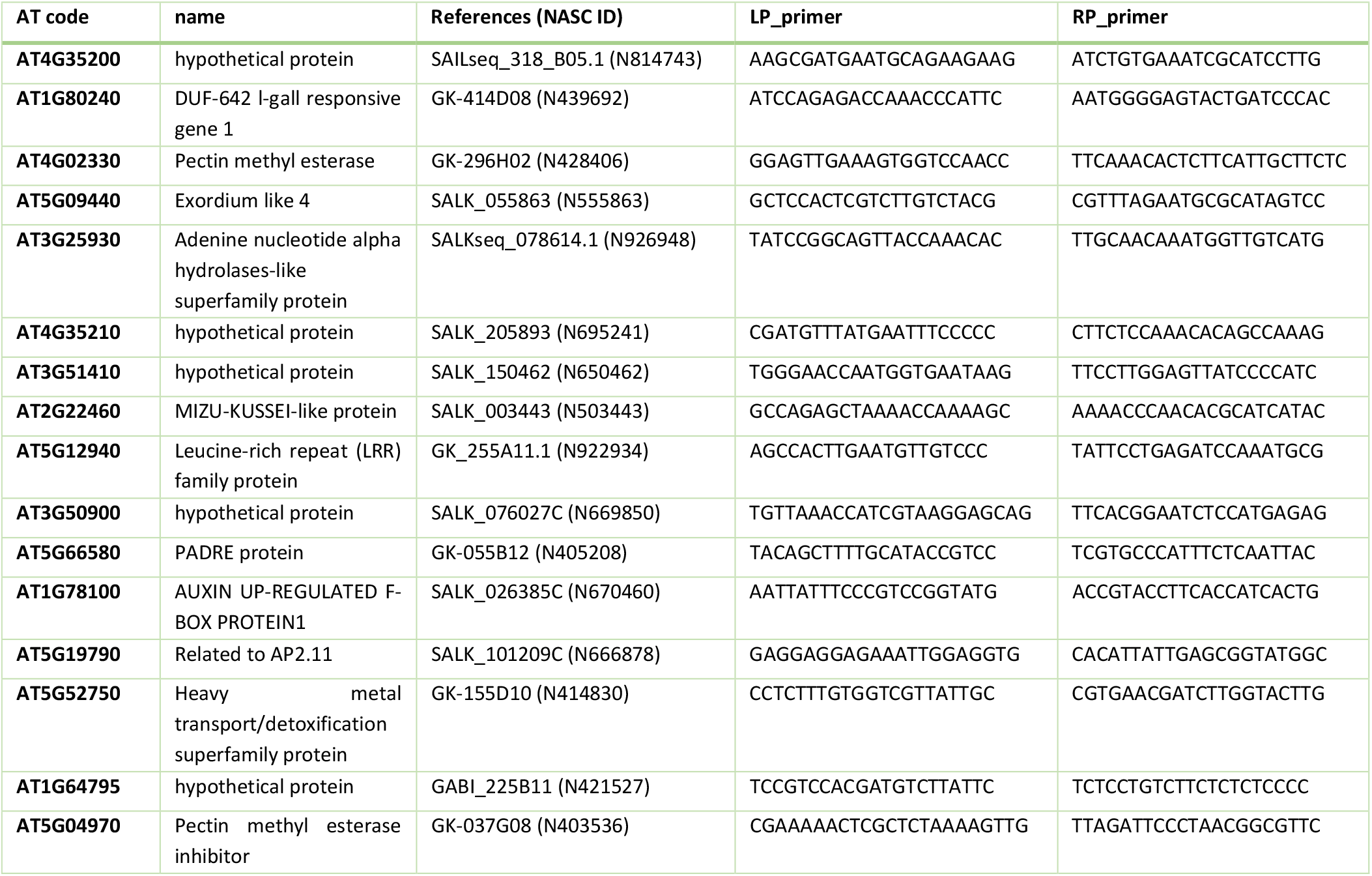
List of mutants and primers used for experiments given in Table S1.

